# Gap junctions set the speed and nucleation rate of stage I retinal waves

**DOI:** 10.1101/368019

**Authors:** Kähne Malte, Rüdiger Sten, Kihara Alexandre, Lindner Benjamin

## Abstract

Spontaneous waves in the developing retina are essential in the formation of the retinotopic mapping in the visual system. From experiments in rabbits, it is known that the earliest type of retinal waves (stage I) is nucleated spontaneously, propagates at a speed of 451±91 *μ*m/sec and relies on gap junction coupling between ganglion cells. Because gap junctions (electrical synapses) have short integration times, it has been argued that they cannot set the low speed of stage I retinal waves. Here, we present a theoretical study of a two-dimensional neural network of the ganglion cell layer with gap junction coupling and intrinsic noise. We demonstrate that this model can explain observed nucleation rates as well as the comparatively slow propagation speed of the waves. From the interaction between two coupled neurons, we estimate the wave speed in the model network. Furthermore, using simulations of small networks of neurons (N≤260), we estimate the nucleation rate in form of an Arrhenius escape rate. These results allow for informed simulations of a realistically sized network, yielding values of the gap junction coupling and the intrinsic noise level that are in a physiologically plausible range.

**Author summary:** Retinal waves are a prominent example of spontaneous activity that is observed in neuronal systems of many different species during development. Spatio-temporally correlated bursts travel across the retina at a few hundred *μ*m/sec to facilitate the maturation of the underlying neuronal circuits. Even at the earliest stage, in which the network merely consists of ganglion cells coupled by electric synapses (gap junctions), it is unclear which mechanisms are responsible for wave nucleation and transmission speed. We propose a model of gap-junction coupled noisy neurons, in which waves emerge from rare stochastic fluctuations in single cells and the wave’s transmission speed is set by the latency of the burst onset in response to gap-junction currents between neighboring cells.

## 1 Introduction

Spontaneous activity spreads through neuronal systems of many different mammal species during development. Crucial roles are attributed to this spontaneous activity [1]. Among the most prominent roles is the synaptic refinement in the retina, where spatio-temporally correlated bursts of activity are observed, and it was found that blocking these waves disrupts eye-specific segregation into the visual thalamus [2, 3]. Therefore, much effort has been devoted in recent years (e.g. [4–7]) to understand the responsible mechanisms of retinal waves. The observed patterns of spontaneous activity in the developing retina are remarkably similar across many species [1]. These patterns have been characterized as spatially correlated bursts of activity in the ganglion cell (GC) layer, which are followed by periods of silence [8–10].

Retinal waves are mediated by three distinct circuits at different developmental stages that have been described in rodents, (for review see e.g. [1]). In stage I (E17-P1), bursts of activity spread between retinal ganglion cells. In this stage, few synapses are identifiable and waves are mediated by gap junctions (GJs) and adenosine [11]. Stage II waves begin with the onset of synaptogenesis and end with the maturation of glutamatergic circuits while stage III waves end with eyeopening and the onset of vision [12, 13]. Here, we exclusively focus on stage I waves, observed prior to the emergence of functional chemical synapses in the retina. These waves show random initiation sites, no directional bias, and a propagation speed of about 450 *μ*m/s. Via patch-clamp recordings, stage I retinal waves were found to be initiated and propagated in the GC layer [11].

In this work we develop a theoretical model of the retina and limit ourselves to a GC layer of bursting neurons which are diffusively coupled by GJs. These electrical synapses are formed between each of the major neuron types in the vertebrate retina [14–18] and play a major role in signal processing and transmission of visual information (for a review, see [18]). GJs are formed by two apposed hemichannels, each one formed by an hexameric array of proteins know as connexins. In mammals, connexin-36 and connexin-45 were clearly identified in neurons located in the inner retina [15, 19]. Both types of connexins follow a distinct expression pattern during retinal development [20]. However, their involvement in the maturing process of the retina is not yet fully understood [21]. GJs have been proposed as the responsible mediator of stage I retinal waves but not yet been used in a model of such waves [5], which is the gap that we intend to fill with our study.

While there is convincing experimental evidence that stage I retinal waves are mediated by GJs, thus far they have not been explicitly addressed with theoretical approaches. (GJ coupling between neurons has been studied theoretically before, e.g. [22, 23]). From a physical perspective, GJs are electrical synapses acting with integration times of the order of milliseconds and were thus argued not to be the mediator of stage I waves [5, 9], which are much slower compared to this time-scale. In this work, we present a model of stage I retinal waves, formed by a network of bursting cells, which are coupled by instantaneously acting diffusive GJs. We show that under certain conditions, the wave propagation can be sufficiently slow to be the responsible mediator for stage I retinal waves. We discuss analytical estimations of the propagation velocities and wave nucleation rate.

## 2 Methods

### 2.1 Model for the Single Retinal Ganglion Cell

We use the phenomenological Izhikevich neuron model, known for displaying biologically plausible dynamics. It similar to conductance-based neuron models of the Hodgkin-Huxley type, while being very efficient from a computational point of view [24, 25]. The model can be regarded as a quadratic integrate-and-fire neuron for the membrane voltage *V_i_*(*t*) of the *i*th neuron with an additional slow recovery variable *u_i_*(*t*), also referred to as gating variable (cf. Fig. 1(a) for the nullclines of the system):

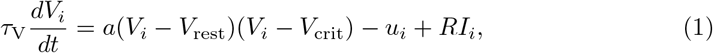

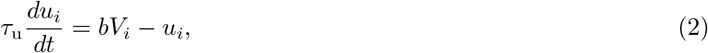

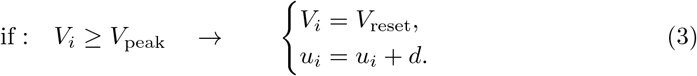

**Fig 1.**
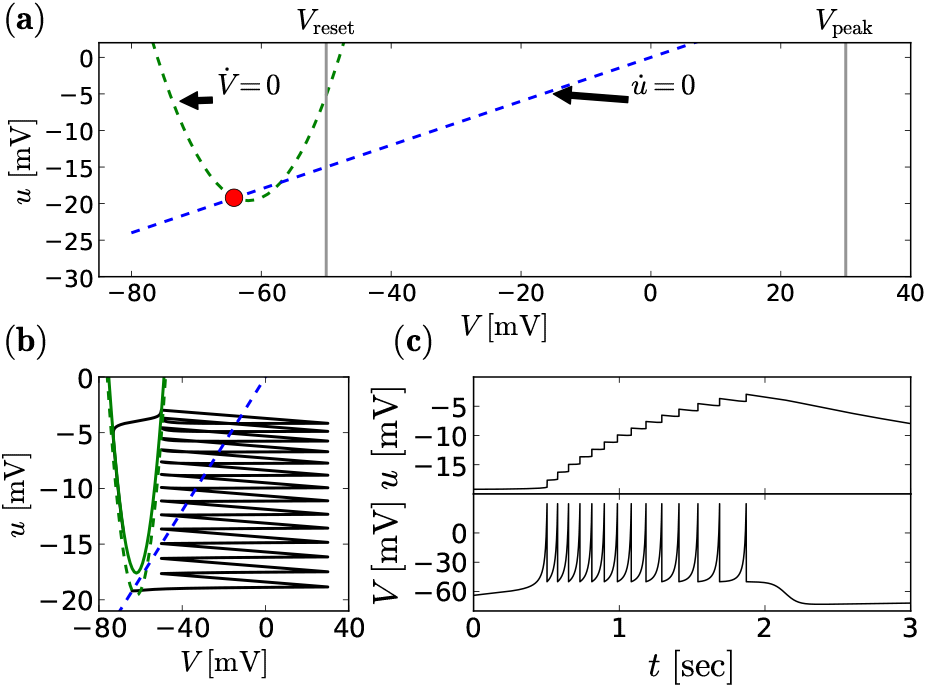
Burst mechanism of the single neuron model. (**a**) shows the nullclines of the Izhikevich neuron model in phase space (*V, u*) without current, *RI* = 0. The green dashed line shows the voltage nullcline and the blue dashed line shows the gating variable nullcline, respectively. Intersections of these two lines are fixed points of the system. The lower fixed point, indicated in red, is stable and represents the resting state of the neuron at (*V, u*) = (*V*_r_*, u*_r_) = (−64mV, −19.4mV). The gray vertical lines indicate the peak voltage *V*_peak_ and the reset voltage *V*_reset_. (**b**) shows the path in phase space of a neuron that is initially in the resting position, but exposed to an external current with *RI* = 2 mV from *t* = 0. The temporal evolution of the separate components *u* and *V* is illustrated in (**c**).

The membrane recovery variable provides negative feedback to the voltage (cf. Fig. 1(b) and Fig. 1(c) top). The parameters *a, b, d* as well as *V*_rest_, *V*_crit_, *V*_reset_, and *V*_peak_ determine the spiking regime of the neuron, with *V*_rest_ *< V*_crit_ *< V*_peak_. The time-scales of the voltage and gating variable are defined by *τ*_V_ and *τ*_u_, respectively. For *u*(*t*) ≡ 0 and *I*(*t*) ≡ 0, *V*_rest_ and *V*_crit_ are the stable and the unstable fixed points of the dynamics, respectively. If *V_i_ ≥ V*_peak_, the membrane potential is reset to *V*_reset_, the *k*th spike time, *t_i,k_*, is registered, and the recovery variable is increased by the constant value *d*. We choose parameters such that the burst characteristics of our model neuron illustrated in Fig. 1 roughly agree with experimental measurements from Syed et al. [11]. Specifically, we aim at a burst duration of about 1 – 2 seconds (cf. Fig. 1(c) bottom) and a spike frequency during bursts of about 5 – 15 Hz. We find those characteristics reasonably met for: *a* = 0.1, *b* = 0.3, *d* = 1.2, *τ*_V_ = 100 ms, *τ*_u_ = 0.0003^−1^ ms, *V*_rest_ = −76 mV, *V*_crit_ = −48 mV, *V*_peak_ = 30 mV, *V*_reset_ = −50 mV. The bursting mechanism is illustrated in Fig. 1.

The total current *RI_i_* = *R*[*I*_gap*,i*_ + *I*_noise*,i*_] is a superposition of the intrinsic noise current and GJ currents from neighboring cells (see below). The intrinsic noise originates from fluctuations of the various channel populations (sodium, calcium, and different potassium channels, see e.g. [26]) and is approximated by white Gaussian noise: 
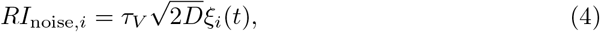
with 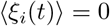and 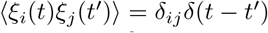and *D* is the noise intensity. We perform simulations at discrete times with a time step of Δ*t* = 0.1 msec according to an Euler-Maruyama integration scheme, see Appendix Sec. 5.

### 2.2 Retinal Network

Ganglion cells are distributed within the ganglion cell layer with a decreasing density towards the outer regions of the retina. For instance, the density in rabbits covers a range from 5000 cells/mm^2^ down to 200 cells/mm^2^ (the mean value is 800) [27]. In a previous study of retinal waves observed in rats, Butts et al. [4] used a ganglion cell density of ∼ 4000 cells/mm^2^. In their simulations they placed neurons in a regular triangular lattice for which the given density translates to a lattice spacing of 17 *μ*m. Because we focus on the rabbit retina, we assume a triangular lattice with a different lattice spacing of 38 *μ*m, reflecting the lower cell density (800 cells/mm^2^) for this system. The reported experimental observations on characteristics of stage I retinal wave were obtained from retina patches of roughly 3 × 5 mm. A mean cell density of 800 cells/mm^2^ translates to a total cell number estimate of 12,000 cells in the studied system. For comparability, we use a similar number of cells for simulations (i.e. 12,100 = 110 × 110).

Here, we ignore for simplicity the inhomogeneous and irregular structure of the ganglion cell layer. We place *N* = *n* × *n* single ganglion cells in a rectangular domain on a triangular lattice such that every cell is connected with GJs to six nearest neighbors, see Fig. 2(c). For illustrative purposes, we will also consider a one-dimensional chain, in which each neuron has only two neighbors. Because we are interested only in stage I waves, prior to synaptogenesis, these cells are not connected to any other cells, i.e. bipolar and amacrine cells are not part of our model. We choose a common approach (e.g. [22]) to model the GJ current as diffusive and instantaneous coupling by 
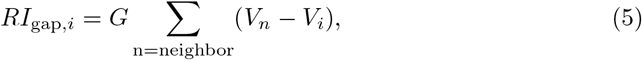
where *G* is the rescaled dimensionless GJ coupling, i.e. *G* = *R*/*R*_gap_. The membrane resistance *R* of retinal ganglion cells can experimentally be measured and is in the range of 100–500 MΩ, e.g. [28]. *R*_gap_ is the GJ resistance between neighboring ganglion cells in the retina, which depends on the connexin type and the transjunctional voltage difference and is roughly *R*_gap_ ≈ 1GΩ [29, 30]. The values of *R* and *R*_gap_ imply a physiological range for our parameter of *G* ∈ [0.1, 0.5]. Because the time course of the action potential produced by our neuron model is only a coarse approximation of the electrophysiological shape of a spike, the GJ coupling may be stronger or weaker than assumed here. This gives additional justification for choosing a wider range of *G*.

**Fig 2.**
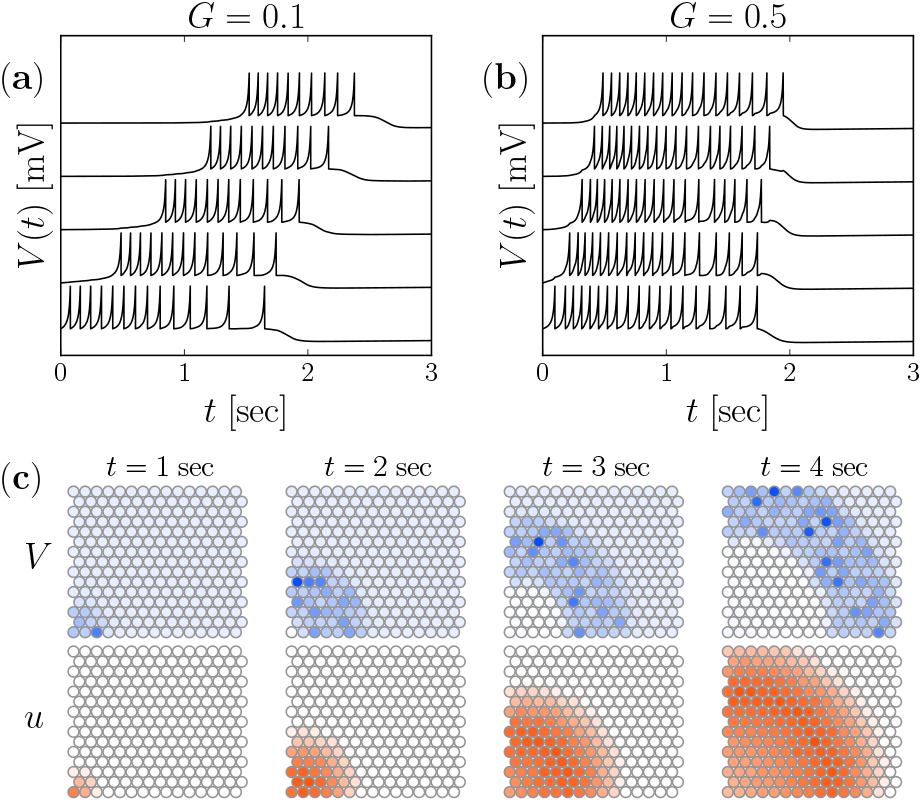
Wave propagation in the deterministic system (*D* = 0). Voltage traces for five model neurons (vertically shifted for better visibility), coupled in a one-dimensional chain with *G* = 0.1 (**a**) and *G* = 0.5 (**b**). The respective first neuron (bottom trace) was initialized in the bursting regime, i.e. (*u*(*t* = 0), *V* (*t* = 0)) = (*u*_rest_, *V*_reset_). Snapshots of waves on a two-dimensional triangular lattice (voltage and recovery variable in top and bottom panels, respectively) with *G* = 0.1 at different time instances as indicated (**c**).

For the two-dimensional setup, we apply two different boundary conditions. For estimating the noise dependence of propagation velocities and nucleation rates, we perform small system simulations (N∼50–260) with periodic boundary conditions in both directions (system on a torus) in order to avoid strong finite-size effects. Simulations of the full system with N∼12,000 are carried out with two additional layers of neurons on the boundary, that are not exposed to intrinsic noise (cells on the system boundary have fewer neighbors, between 2 and 5 instead of 6). Neurons in the two outer layers of the large simulations are discarded from all statistical evaluations.

Single propagating waves running through the network can be captured by the population activity [31] 
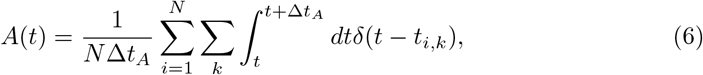
which is a firing rate that is averaged over the network and over a time-bin Δ*t_A_*. We use Δ*t_A_* = 0.5 seconds, which is comparatively large and covers multiple spikes when the cells are bursting.

## 3 Results

### 3.1 Wave Propagation

If we couple cells in a chain and initiate a burst in one of them, we see a propagation of the burst along the chain (Fig. 2(a)). A higher propagation speed can be achieved by increasing the GJ conductance parameter *G* Fig. 2(b). The picture is similar in our two-dimensional setup, for which snapshots are shown in Fig. 2(c). In this case, the wave has been evoked by enforcing a burst in the lower left corner. It propagates as a circularly shaped wave front, which is a consequence of the regularity and rotational symmetry of the system. The gating variable *u* (lower row in Fig. 2(c)) can be associated with the experimentally accessible calcium dynamics and resembles calcium fluorescences images [11]. Compared to the membrane potential (top row), the wavefront of the gating variable lags behind, as it slowly builds up during the burst.

In both, one-dimensional and two-dimensional simulations in Fig. 2, we have set the intrinsic noise intensity to zero in order to illustrate that wave propagation does not hinge on the presence of fluctuations. We note already here, that the propagation speed in the two-dimensional system matches the order of magnitude of biologically observed values. To determine the speed of the waves from simulation such as shown in Fig. 2(c), we approximate the wave’s shape as circular with a fixed center. We define a wavefront as the group of neurons that spike within the same time bin of Δ*t* = 0.1 seconds (see left illustration in Fig. 3(a)) and measure the front’s mean distance from the center and its mean time instance of occurrence. From the differences of these distances and times, we determine the mean velocity, which we find to be weakly distance dependent, but saturating at about 350 *μ*m from the origin of the wave, cf. Fig. 3(b). In the following, all velocity values are averaged over measurements for the range of distances 350 – 650 *μ*m (shaded area in Fig. 3(b)) from the point of initiation and we refer to this measuring method as concentric method. The velocities are shown in Fig. 3(c) as a function of the GJ parameter for the physiologically relevant range of *G* (see methods). We obtain velocities that are in the range of values observed in the rabbit retina [11], cf. the shaded area in Fig. 3(b). The experimental mean value of about 450 *μ*m/sec is attained for *G* ≈ 0.4.

**Fig 3.**
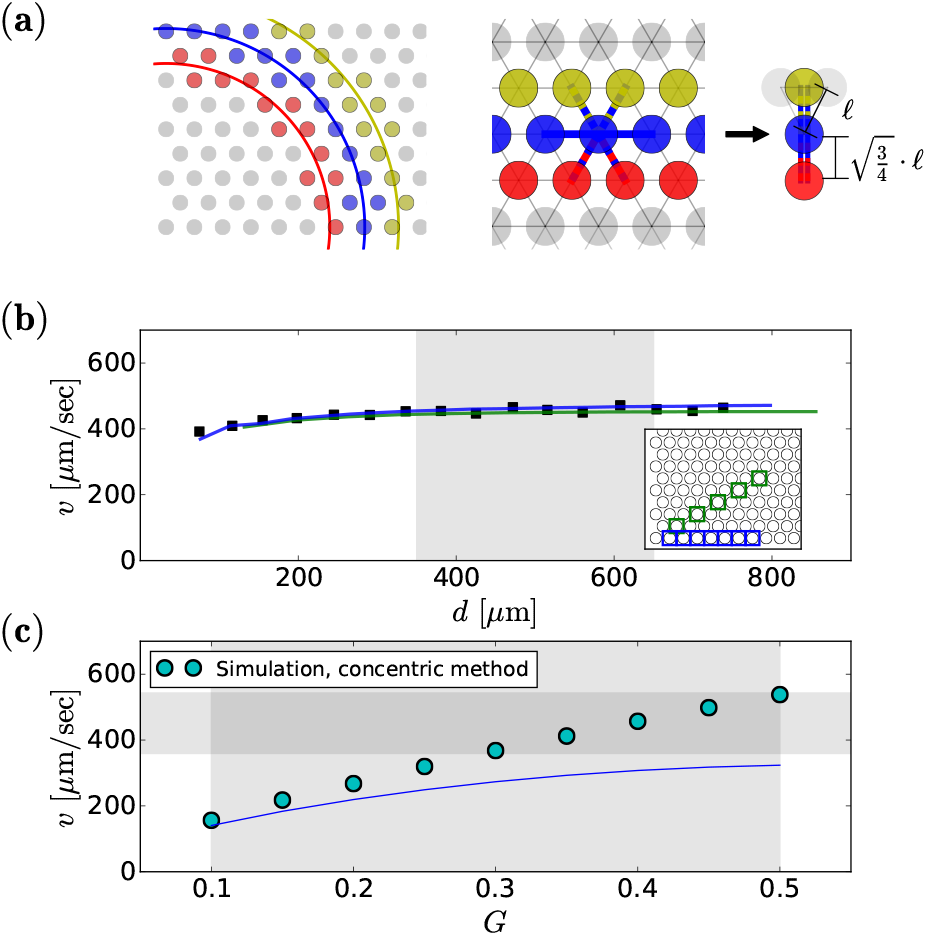
Wave speed: measurement and dependence on GJ coupling. Neural groups with simultaneous burst onset of an exemplary simulation (time resolution Δ*t* = 0.1 seconds) are shown in the panel (**a**) left, for three consecutive time bins in different colors. At large distances from the origin, the shape of a wavefront can be approximated as planar, cf. (**a**) middle. The mechanism of burst propagation can then be mimicked by a one-dimensional situation. Therefore in our theoretical derivations, the distance and coupling strength has to be modified, cf. (**a**) right and details in the main text. Squares in (**b**) represent the speed of the concentric wave (*G* = 0.4) as a function of the distance from the wave’s origin (lower left corner of the simulation domain), measured as described in the text. Alternatively, the speed can be assessed by measuring burst onset times along different fixed directions of the network, i.e. at blue and green sites shown in the inset of (**b**). The resulting wave speeds as functions of distance (blue and green lines) agree closely with the concentric method (squares in (**b**)). The speed shown in (**c**) is the mean value of simulation data (symbols) of the shaded area in (**a**) as a function of the GJ coupling *G*. Simulation results are compared to *v*_2*D*_(*G*), eq. (9). The vertical and horizontal shadings indicate the physiological range of *G* (see methods) and the observed velocities in the rabbit retina [11], respectively.

The propagation and its speed can be theoretically understood as follows. Assuming a steep wave profile, the speed of the wave is given by the inverse of the time it takes a bursting neuron to excite its neighbors, times the displacement of the corresponding wave fronts. We refer to this time as burst onset time difference (BOTD). For simplicity, we neglect noise and consider in the following a one-dimensional setup consisting of three neurons: one initially quiescent neuron (*i*) is connected to a bursting neuron (*i −* 1) on one side and to a quiescent neuron (*i* + 1) on the other side. They are separated by the lattice spacing *ℓ* = 38 *μ*m, hence the velocity is defined as *v*_1D_ = *ℓ*/*T*_B_. Therein, *T*_B_ denotes the analytical approximation of the BOTD for this one-dimensional case.

The approximation *T*_B_ for the BOTD between neighboring neurons can be derived using three assumptions (details in the Appendix, Sec. 5). First, we assume a constant gating variable (*u*(*t*) ≈ *u*_r_ = const), which is reasonable on a short time scale, because τ_u_ ≫ τ_V_. Second, we replace the voltage variable of the bursting neuron *V_i–_*_1_(*t*) by its temporal average 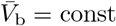, that can be analytically calculated (see Appendix) and for our standard parameters is 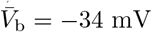. Third, we replace the voltage of the quiescent neuron that is not directly connected to the bursting neuron by the resting potential, *V_i_*_+1_ = *V*_r_. Consequently, the GJ current seen by the driven neuron reads 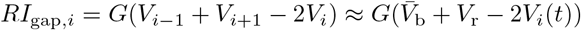, and the resulting dynamics until the voltage *V_i_* reaches the peak potential for the first time is effectively one-dimensional and can be recast to the form (refer to Sec. 5 for details):
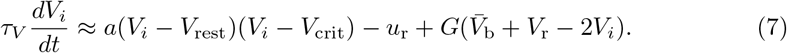

This first order ordinary differential equation can be solved via separation of variables to find *t*(*V*). We obtain it by first calculating the difference of the times from the voltage being at its peak potential and its resting potential. However, the driven neuron is already exposed to the driving GJ current while the voltage of the bursting neuron travels to its first spike time. Therefore, for simplicity we subtract the first inter-spike interval *T*_ISI_ from the beforehand calculated time difference: 
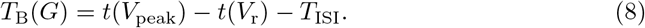

The explicit expression is lengthy and derived in the appendix (resulting in eq. (24)). Comparing *T*_B_ to simulations of a one-dimensional chain shows a reasonable agreement (see Appendix), although the theory overestimates the simulated values, in particular, for larger values of *G*.

In the two-dimensional setup at larger times, the wave attains a planar shape as indicated in Fig. 3(a), where red circles represent bursting neurons and blue and yellow circles represent driven and quiescent neurons. Now, we assume that the wave front is perfectly flat and all neurons shown in the same color share an identical voltage. In that case, the propagation mechanism simplifies to two bursting neurons exciting one quiescent neuron, whose membrane potential is further affected by two quiescent neurons. Hence, we can mimic the quasi one-dimensional situation by doubling the value of *G* and additionally taking into account the modification of the effective length, i.e. *ℓ*_eff_ = (3/4)^1/2^*ℓ*, see Fig. 3(a). Consequently, we can approximate the velocity in the two-dimensional system as 
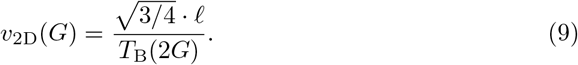

Calculated velocities *v*_2D_(*G*) are shown in Fig. 3(c) by the blue line, underestimating the true velocity (circles) but providing a correct order-of-magnitude estimate. Note that so far we restricted the considerations to a purely deterministic setup. Our Simulation, concentric method simulations with noise indicate that moderate fluctuations have only little impact on the mean velocities.

### 3.2 Wave Nucleation

In the stochastic version of our system, we observe spontaneous waves that resemble those found in experiments [11]. Experimentally, it was observed by Syed et al. [11] that the spontaneously nucleated waves appear with a mean inter-wave interval *T*_IWI_ of 36 seconds. In our model, waves are initiated by noise, since neurons are set in the excitable regime and cannot generate periodic spiking or bursting without external input. We expect that the nucleation rate per neuron depends strongly on the noise intensity *D*. To characterize this dependence, we simulate small systems (N∼50–260, see methods) with periodic boundary conditions for two different values of GJ coupling and different noise intensities, cf. Fig. 4.

**Fig 4.**
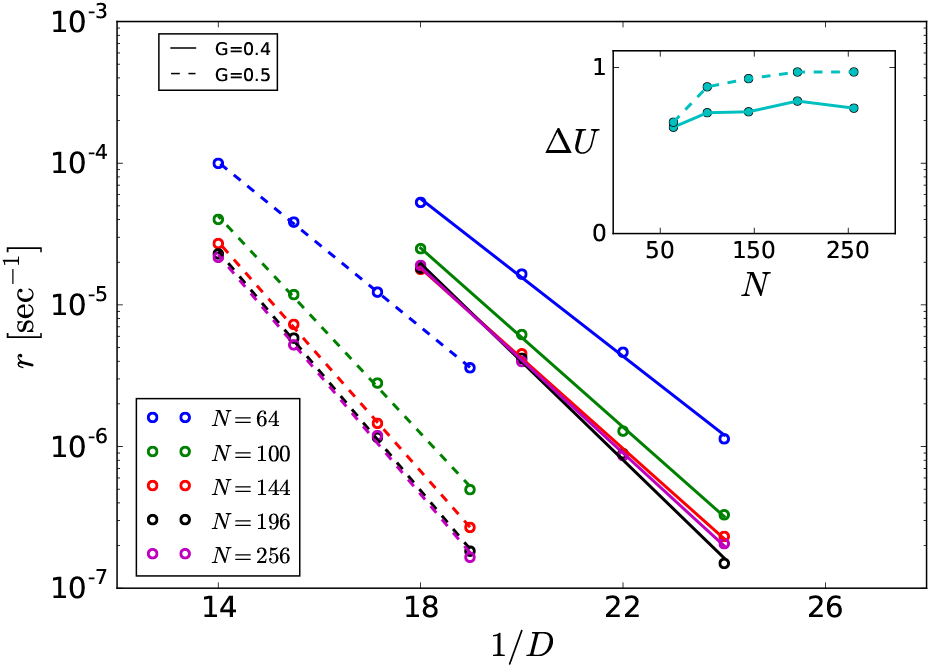
Arrhenius Plot. Spontaneous nucleation rate as function of the inverse noise intensity obtained from four two-dimensional systems with different system sizes as indicated and periodic boundary conditions. From the linear fit of these data, an effective potential barrier *ΔU* and a rate prefactor *r*_0_ can be estimated (dependence of *ΔU* on system size shown in inset).

With the understanding that every neuron has the same chance to trigger a wave, the global nucleation rate should be linear with *N* to a first approximation. Thus we measure the nucleation rate per neuron as *r* = 1*/*(*T*_IWI_*N*). As demonstrated in Fig. 4 by the linear dependence of the rate’s logarithm on the inverse noise intensity, we obtain an Arrhenius rate 
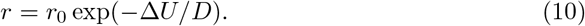

The effective potential barrier Δ*U* depends on *G* and the system size *N* and saturates for sufficiently large systems (inset) for both values of *G*.

The increase of the potential barrier with *G* can be understood to first approximation by the effective change of the current-voltage relation in the single neuron. The GJ coupling term eq. (5) leads to an effective increase in the leak current that stabilizes the resting potential and makes it harder to initiate a burst. This mechanism is dominant in comparison to the influence of other coupling effects and the stochasticity of the neighbors on the nucleation rate (supported by additional simulations, see appendix).

The more subtle dependence of Δ*U* on the system size can be explained as follows: Coupling stochastic neurons in small systems with periodic boundary conditions leads to spatial correlations and thus effectively to stronger noise. This effect can be neglected for large system sizes or weak coupling, but has a measurable effect otherwise (cf. Fig. 4 and Fig. 4 inset).

### 3.3 Discussion of Large-Scale Simulation Results

Our results so far can be used to predict the mean inter-wave interval and the propagation speed of retinal waves for a system size *N* = 12,100 that roughly corresponds to the experimentally studied patch size in Ref. [11]. Vice versa, we can infer an approximate value of the noise intensity *D* that leads to the experimentally observed value of *T*_IWI_ = 36 seconds and test this in numerical simulations of the full system.

For our estimation of the rough value of the noise intensity in a large system, we have to take into account that the single neuron undergoes a substantial refractory period of *T*_ref_ ≈ 14 seconds after bursting (estimated from small-system simulations investigating the minimal inter-wave interval for various noise intensities). The mean inter-wave interval is then given by *T*_IWI_ = *T*_ref_ + 1*/*[*N · r*(*D*)] and the estimated value of the noise intensity follows from the Arrhenius law, eq. (10), as 
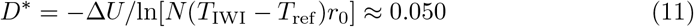
(for *G* = 0.4, and *r*_0_ = 6 and Δ*U* = 0.71, fit parameters from Fig. 4, solid line with *N* = 256).

The estimated parameters, *G* = 0.4 and *D* = 0.050, can now be used in a large-scale simulation. In Fig. 5(a), we show snapshots of the full system’s gating variable (a proxy for the experimentally accessible calcium concentration). The wave front seen in the experimentally observable area (box in Fig. 5(a)) looks similar to experimental measurements, cf. Ref. [11]. From Fig. 5(b), it becomes evident that the mean inter-wave interval becomes much shorter for a slight increase in *D*. The mean inter-wave interval at these parameter values is not exactly 36 seconds, but somewhat larger: these statistics depend very sensitively on the value of the noise intensity (i.e. on the second leading digit, cf. Fig. 5(c) middle). This is seen in the global population activity, that reveals a wave going through the system as a single peak vs. time.

**Fig 5.**
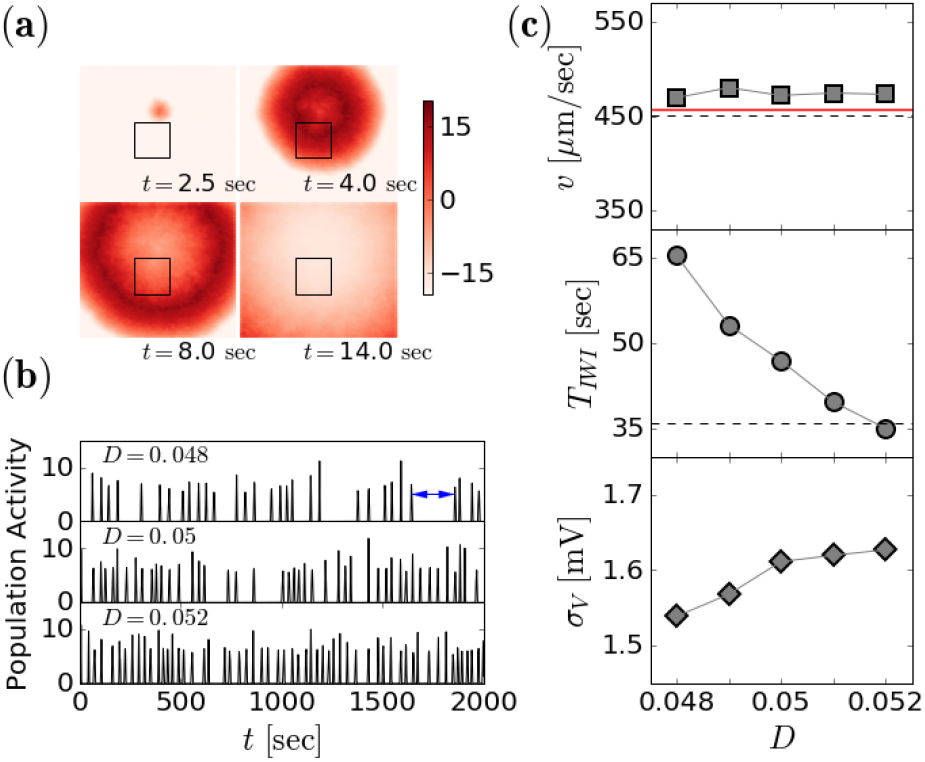
Large-Scale Simulations. A network of 12,100 GJ coupled and noisy Izhikevich neurons display spontaneously nucleated waves that propagate with velocities the are comparable to experimentally observed values. (**a**) Snapshots of the gating variable (associated to a proxy for the calcium concentration) at different time instances during one wave running through the system (*G* = 0.4, *D* = 0.05). The small rectangle indicates the dimensions of the experimentally accessible observation area [11]. (**b**) Population activity *A*(*t*) (with Δ*t*_A_ = 0.5 seconds, see eq. (6)) of the entire system over a larger time window for different noise levels. One wave, as shown in (**a**) collapses here into a single peak; time differences between adjacent peaks are the inter-wave intervals *T*_IWI, i_ (one indicated by an arrow). (**c**) Mean velocity, mean inter-wave interval and standard deviation of the subthreshold membrane voltage as a function of the noise intensity for a small range around the estimated value *D*_target_ = 0.05. Dashed black lines indicate experimental mean values from Ref. [11], solid red line shows the wave speed for *D* = 0, extracted from the circle at *G* = 0.4 in Fig. 3.

The dependence of crucial neural statistics on the noise intensity is illustrated in Fig. 5(c). In contrast to the mean inter-wave interval, the mean velocity of the wave does not depend strongly on the noise (Fig. 5(c), top) but stays close to the experimentally observed mean value (dashed line). The fine tuning of the noise intensity shows that the experimental value of 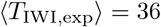 seconds is attained for a noise level of *D* = 0.052, slightly larger than *D^*^* (estimated above). How realistic is this noise level? To address this question, we show at the bottom of Fig. 5(c) the standard deviation of the subthreshold voltage fluctuations, σ_V_, as a function of the noise intensity *D*. σ_V_ increases only slightly with *D* and attains values around 1.6 mV.

To our knowledge, there are no detailed investigations of intrinsic noise sources in retinal ganglion cells at embryonic age. Because in this developmental stage there are no chemical synapses present [32], the synaptic background fluctuations can be excluded for our system. One likely source of variability is channel noise that typically leads to small membrane potential fluctuations with a standard deviation σ_V_ below 0.6 mV [33, 34]. The noise intensity that is required for the experimentally observed inter-wave interval results in sub-threshold voltage fluctuations that are three times bigger, cf. Fig. 5(c) bottom, suggesting that besides ion channel noise there are additional sources of fluctuations present. These could result from stochasticity of GJs itself but also indirectly from GJs via differences in individual resting potentials (for the heterogeneity of the resting potential in similarly sized cells, pyramidal cells in the cortex, see [35]). In any case, the apparent voltage fluctuations of about 1.6 mV are well within the range of experimentally observed voltage noise in embryonic ganglion cells (cf. Fig. 1 in Ref. [11]).

## 4 Summary

The investigations presented in this paper propose a GJ-based model of stage I waves in the developing retina. Starting with a neuron model that roughly reproduces the spiking properties of a burst of one single retinal ganglion cell, we incorporated GJ coupling of physiologically plausible strength and temporally uncorrelated fluctuation. This allowed us to reproduce the characteristics of wave nucleation and slow wave propagation in the early retina. Earlier it was believed that GJs can play a role in fast neural transmissions only [5, 9], since the current in electrical synapses responds much quicker than neurotransmitters in chemical synapses. As shown in our paper, however, it is possible to obtain a limited transmission speed in a simple Ohmic model of the GJ coupling. Furthermore, although stochastic fluctuations are strong enough to ignite bursts with the correct nucleation rate, they do not distort the propagating fronts very much, i.e. the wave propagation is still a reliable process.

The reason for the slow transmission we observe can be found in the nonlinear dynamics of the single neuron. The Izhikevich model that we use for the ganglion cell is essentially a quadratic integrate-and-fire neuron model with a slow adaptation variable. This model is the normal form of a saddle-node bifurcation and has a pronounced latency if close to this bifurcation, i.e. the spike response to a current step (in our case provided by a neighboring bursting cell) is considerably delayed because the system experiences the ”ghost of the former fixed point”, see Ref. [36]. The presence of weak noise modifies this picture only slightly [37].

Although our model accounts for the most important features of wave nucleation and propagation for stage I retinal waves, it cannot explain the strong variability of the experimentally measured statistics (error of velocity ±91 *μ*m/sec [11]). This is due to a number of model simplifications, which we now concludingly discuss. Firstly, the real system is much more heterogeneous than in our model; secondly, GJs may couple more than next neighbors and their conductivity may be noisy and voltage gated; thirdly, the detailed dynamics of ganglion cells is certainly more complex than can be captured by the Ihzikevich model; last but not least, the white Gaussian noise in our model is a rather coarse approximation of the channel noise and other fluctuations in the system.

In our model, we arranged the neurons on a highly regular lattice with a cellular spacing according to an experimentally determined mean value of cell density, neglecting the strong heterogeneities in the distribution [27]. On this lattice, each cell is connected to exactly six nearest neighbors. Given the aforementioned heterogeneity, the numbers and distances between neighbors will be more broadly distributed than in our model. Incorporating these heterogeneous features in the simulations would likely broaden the range of observed velocities and thus better reflect the considerable variability found in experimentally measured values.

The soma size of (rabbit) retinal ganglion cells (< 30*μ*m, e.g. Ref [27]) is smaller than our employed lattice spacing, implying GJ coupling between dendrites rather than soma-soma coupling only. The size of the dendritic arbor of retinal ganglion cells is ∼ 100 − 130*μ*m, thus suggesting direct communication between cells that are up to the threefold of the lattice spacing apart. In our simulations with only next-neighbor coupling, we could reproduce the experimentally observed velocity with a comparatively large coupling constant of *G* = 0.4 (physiological range was *G* ∈ [0.1, 0.5], see methods). It is conceivable, that this large *G* value is an effective description of a system with larger effective gap-junction neighborhood but with a smaller (and possibly distance-dependent) coupling value *G*. Put differently, we expect similar results for the wave speed in a system with extended coupling neighborhood but reduced coupling strength per connection (with the latter still being within the physiological range).

Regarding the neuron model and the incorporation of noise, we note that for developed retinal ganglion cells detailed multi-compartment conductance-based models with stochastic ion channels exist [26]. With more electrophysiological data available, it will certainly be possible to develop biophysically more realistic models of the bursting ganglion cell at the early stage. Furthermore important for our problem will be the incorporation of stochastic models of GJs [38] with voltage-dependent kinetics [39, 40] and the heterogeneity of physiological parameters such as the resting potential. Such detailed models are certainly difficult to simulate for large networks but could be employed to estimate the total noise intensity in the system and to identify the dominant noise source, cf. similar approaches in Refs. [26, 41, 42].

## 5 Appendix

In the main text we used an analytical approximation of the mean membrane potential during a burst to derive an estimate for the propagation speed of gap junction mediated waves. In the following, we provide all details necessary to arrive at the equations that we discussed above.

## 5.1 Approximate solution of the voltage equation with constant parameters

All approximations are based on solving the time dependence of the membrane potential, eq. (1), which becomes analytically feasible only by decoupling the system Eq. (1–3). We are specifically interested in the propagation mechanism, and thus in the time the voltage needs to go from the vicinity of the resting potential to the peak potential that marks the occurrence of the first spike. Consequently, the reset mechanism eq. (3) can be discarded. Until this first spike time, the gating variable *u* can roughly be regarded as constant, cf. the phase space trajectory displayed in Fig. 1(b). Accordingly, we replace *u_i_* in eq. (1) by the constant *u*_r_ = −19.2 mV., i.e. the resting value of the gating variable. The resulting expression is still a system of coupled differential equations, due to the interaction with neighboring neurons in the network, that is given by the gap junction current eq. (5). To circumvent this problem we simply replace the membrane potential of a bursting neighboring neuron by the constant value 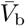, an estimate of the temporal average of a neuron’s voltage during the burst, which is described in detail below. The voltage of the silent neighbors is replaced by the resting potential. For the discussion of the propagation mechanism these replacements are fairly reasonable, because the propagation process can be thought of as a certain number of synchronously bursting neurons (typically, one or two) exciting a connected neighbor with additional quiescent neighbors.

Under these conditions, the dynamics eq. (1) can generally be recast into the form 
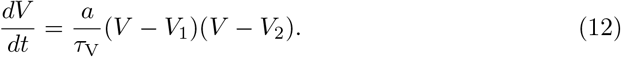

This equation can be solved by separating the variables and integrating. It is however necessary to distinguish the two cases of *V*_1_ and *V*_2_ being real valued (corresponding to the situation with a stable resting potential) or complex valued (unstable situation). We will refer to real valued (*V*_1_*, V*_2_) as (*V*_r_*, V*_c_) with *V*_r_ < *V*_c_. Real values are e.g. obtained for the case of an isolated neuron, or equivalently *G* = 0, and are related to our original parameters as follows 
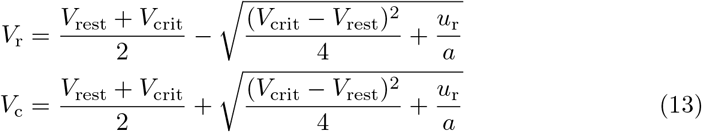
leading to (*V*_r_*, V*_c_) = (−64 mV, −60 mV) for our parameter choice. In this case, the integration of eq. (12) yields 
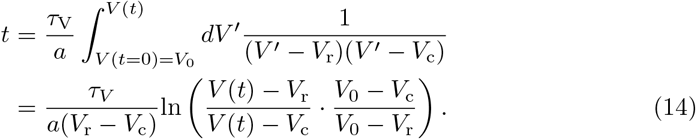
which can be inverted and thus leads to the explicit solution for the voltage trajectory: 
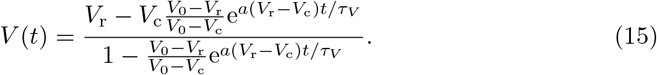

Turning to the case of complex valued (*V*_1_*, V*_2_), we will refer to them as (*V*_m_ + *iγ*, *V*_m_ − *iγ*). Integration of eq. (12) yields

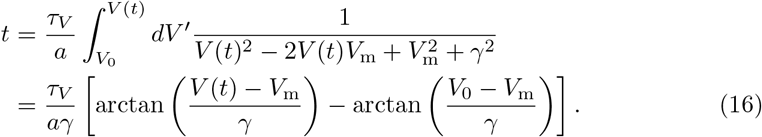

The explicit expression of *V*_m_ and γ depend on the specific setup of the neighbors (see below).

## 5.2 Approximation of the mean voltage during a burst

To analytically understand the propagation of the burst from cell to cell, we approximate the time-dependent driving voltage of the bursting cell by its time average, 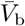, resulting in the GJ current: 
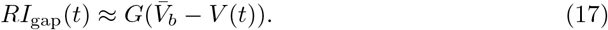

This replacement is justified because during the period of interest (the time needed for the driven cell to generate a spike), the bursting neuron generates at least a few spikes, i.e. changes rapidly compared to the voltage of the driven neuron.

The mean voltage during the burst can be estimated by integrating the voltage over one inter-spike interval *T*_ISI_, for simplicity considered for an isolated cell (*G* = 0) that is started at *V* (*t* = 0) = *V*_reset_, *u*(*t* = 0) = *u*_r_, 
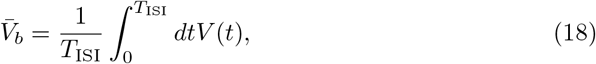
where *T*_ISI_ = *t*(*V*_peak_) − *t*(*V*_reset_). Using eq. (14), we find 
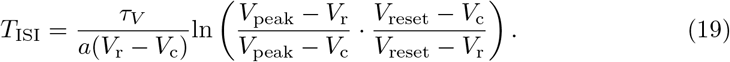

Using eq. (15) and eq. (19) in eq. (18), we can calculate the integral and further simplify the resulting expression 
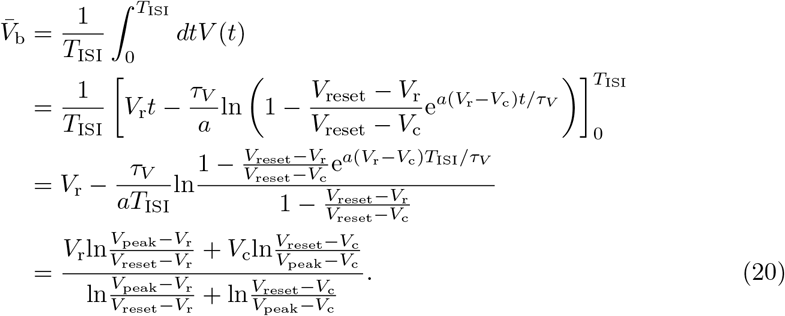

For our standard parameters this gives 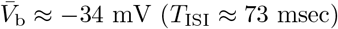.

## 5.3 Approximation of the propagation speed

Let us now turn to the situation, where an initially quiescent neuron is driven to a burst due to the gap junction current that results from one bursting neighbor in a one-dimensional chain. To this end, we consider the three neurons *i −* 1 (bursting), *i* (to be excited), and *i* + 1 (quiescent). Neuron *i* is brought from the resting potential *V*_r_ to the peak potential *V*_peak_ in a period *T*_ISI_ + *T*_B_ that approximately consists of one inter-spike interval *T*_ISI_ (the time needed by neuron *i −* 1 to go from reset to peak potential for the first time) and the BOTD *T*_B_ that determines the speed of the wave via 
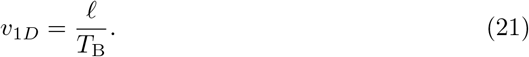

During the period *t* ∈ [−*T, T*_B_] (light blue shaded area in Fig. 6), we approximate the membrane potentials of the bursting and quiescent neurons as 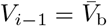 and *V_i_*_+1_ = *V*_r_, respectively, leading to 
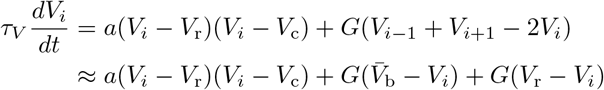

**Fig 6.**
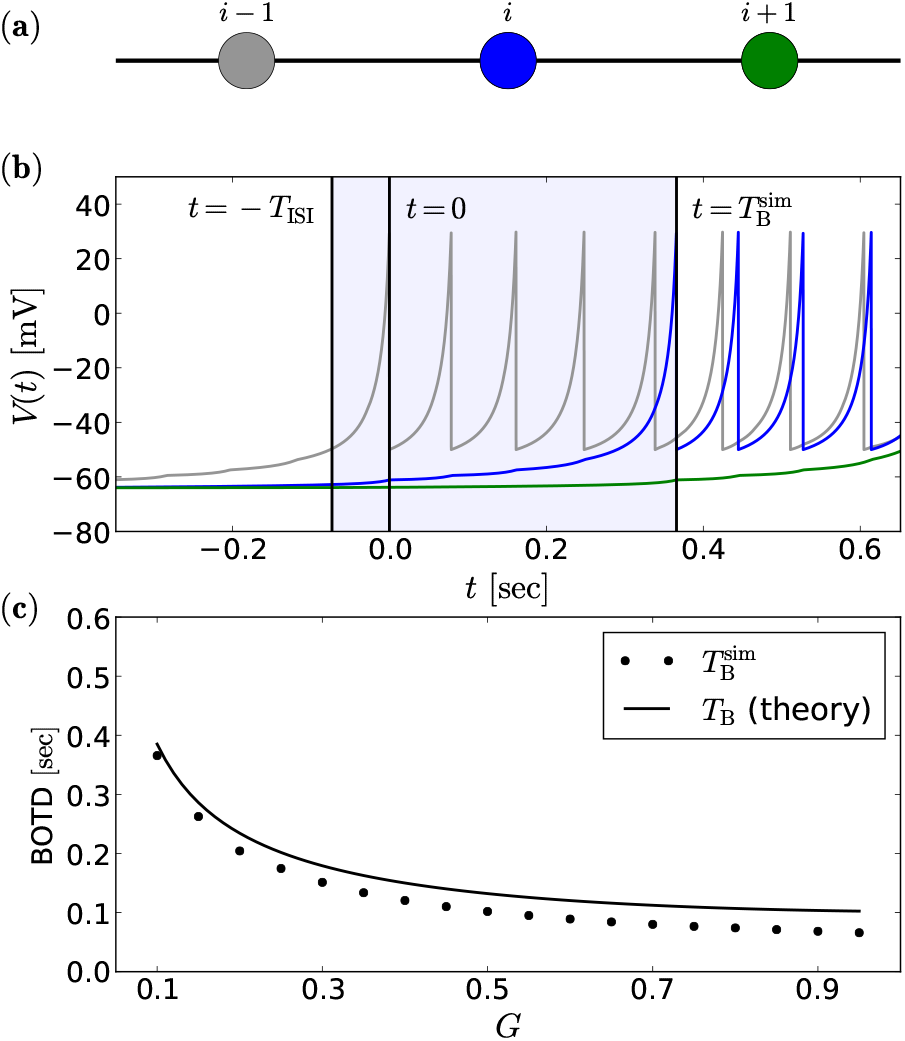
Burst propagation and burst onset time difference in the one-dimensional chain. Schematic illustration of the one-dimensional setup (**a**). Voltage traces of the neurons in (**a**) for *G* = 0.1 shown in (**b**) in respective colors. The shaded area indicates the time period relevant for the excitation of neuron *i* (blue line) by the GJ current from neuron *i* − 1 (gray line). The membrane potential of neuron *i* + 1 (green line) is approximately constant at *V_i_*_+1_ ≈ *V*_r_ during this period. Burst onset time difference as function of *G* is shown in (**c**); simulations (symbols) of a one-dimensional chain without noise (eq. (1–5) with *D* = 0) compared to eq. (24) (solid line).

This equation can be recast into the form of eq. (12) with complex *V*_1, 2_ = *V*_m_ ± *iγ* in the case of propagating bursts, where 
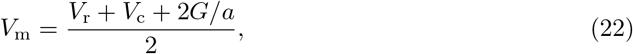
and 
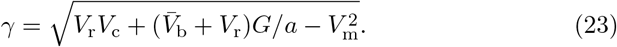

For the calculation of *T*_B_ we can then employ eq. (16), yielding 
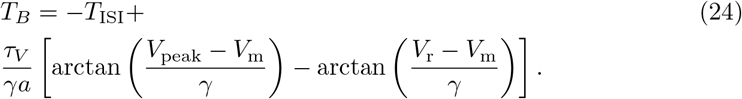

A comparison between calculated burst onset times according to eq. (24) and *T*_B_ obtained from the simulation of a one-dimensional chain is shown in Fig. 6(c). The approximation of *T*_B_ (eq. (24), solid line) shows reasonable agreement with the simulation results (symbols). As stated in the main text, we approximate the propagation speed of waves passing through a two-dimensional network of neurons by eq. (9). This corresponds to the assumption that the wave’s front is reasonably well approximated by a planar shape, if far enough from its origin. As displayed in Fig. 3(b), the propagation mechanism can then be mimicked by a one-dimensional situation with rescaled distance and coupling strength.

In order to see this, note that neurons of the same color in Fig. 3(b) that are part of a perfectly planar wave front share exactly the same state (*V, u*). If the voltage of all horizontal neighbors is identical, links between these neurons can be discarded, because the GJ current is zero. We set the first spike time of the bursts of all red neurons as time origin. Now, every single blue neuron feels an excitatory current from two bursting neurons (connected via the links indicated in red/blue). The leak current of this one blue neuron is affected by the two links connecting this neuron to two yellow neurons (indicated by blue/yellow lines in Fig. 3(b)), which are to a good approximation at rest at this instant of time. Doubling the excitatory current and the additional leak current via gap junctions can be expressed by doubling the GJ conductance parameter *G* in eq. (24). Last but not least, the propagation of the wave within one burst onset time difference is not in direction of the link, but we have to consider the reduced distance 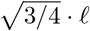. Putting everything together, we obtain eq. (9).

## 5.4 Numerical Simulation Methods

The numerical simulations of our system were performed according to the following Euler-Maruyama integration scheme: 
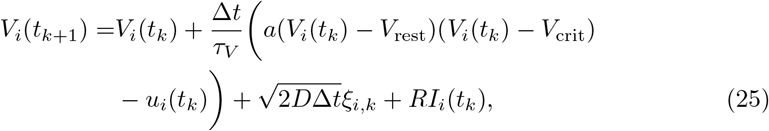

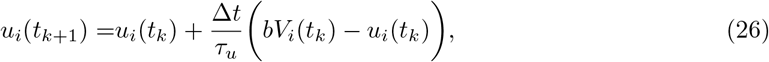
where *ξ_i,k_* are independent Gaussian random numbers with unit variance [43]. The simulation results shown in Fig. 1, Fig. 2, and Fig. 3 are deterministic, i.e. *D* = 0. The wave nucleation rates shown in Fig. 4 correspond to an average of three independent simulations for each set of parameters *G, N*, and *D*, where every single simulation was run until a fixed number of spikes was generated (7500 · *N*), such that a single simulation returned roughly 200 inter-wave intervals; for cases with very low nucleation rates (< 10^−6^), fewer inter-wave intervals were simulated due to a hard-coded time limit. All simulations were performed at discrete times with a step of Δ*t* = 0.1 msec. To test the stability of the Euler-Maruyama integration scheme for our network model for *D* > 0, we compared simulations for the nucleation rate, cf. Fig. 4, with different integration time steps at the set of parameters: *G* = 0.4, *N* = 100, *D* = 1*/*18. We found that reducing the integration time step by a factor 10 had no significant impact on the result of the nucleation rate.

In Fig. 4, we find a finite size effect that vanishes for larger system sizes. The boundary conditions introduce a measurable effect on the spontaneous nucleation rate of retinal waves for smaller system sizes. This effect is due to correlations of distant neighbors in the system. To gain a better understanding of the magnitude of correlation on voltage fluctuations, we simulated a one-dimensional chain of neurons. Because we are interested in the sub-threshold voltage fluctuations we simplified the neuron model to a version that does not generate spikes, i.e. with a linearized deterministic part of the dynamics (GJ and noise current unchanged) at the stable fixed point: 
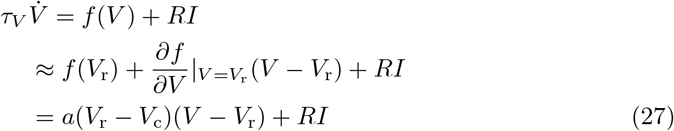

Simulation results for the Pearson correlation coefficient of the membrane potential as function of distance for a ring of 15 coupled neurons are illustrated in Fig. 7(a). With increasing coupling strength, there is a non-zero correlation between the voltages, even for neurons as far as 4 space units apart. This leads to a measurable increase of the overall membrane fluctuations of a ring with chain length up to 5 neurons for *G* = 0.5, cf. Fig. 7(b). For larger chain lengths, the periodicity has no effect on the voltage fluctuations, which is in agreement of the saturation observed for Δ*U* in Fig. 4.

**Fig 7.**
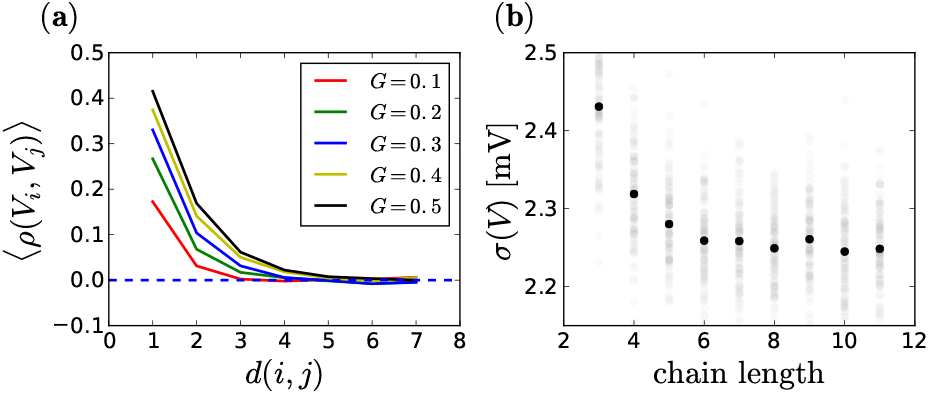
GJ coupling leads to correlations that effect the noise intensity. Pearson correlation coefficient of the voltage as function of the distance between neurons in (**a**). Standard deviation of the membrane potential as function of the chain length for *G* = 0.5 and *D* = 0.05 in (**b**). Each point corresponds to an average of 100 simulations of 1000 seconds for a single neuron (the result of single simulations is illustrated by transparent symbols).

The results for the wave speed shown in Fig. 5 were obtained by averaging the properties of roughly the 20 first waves (by limiting the total number of evaluated spikes to 600 · *N*) from one large scale simulation. The results for the inter-wave intervals were obtained by averaging over all recorded inter-wave intervals of one simulation for every set of parameter. Simulations were run for 5000 seconds with a break criteria at approximately 300 waves (7500 · *N* spikes). Panel (b) of Fig. 5 shows the first 2000 seconds of the population activity for three exemplary noise intensities. Separate simulations were performed to estimate the amplitude of the subthreshold voltage fluctuations; to this end, we used the membrane potential of 20 neurons for 20 seconds, in which no wave was observed.

## Acknowledgments

This work was supported by the Deutsche Forschungsgemeinschaft within the international research training group 1740. We would like to thank Christophe Haynes for their comments on the manuscript.

